# Selfish herd effects in aggregated caterpillars and their interaction with warning signals

**DOI:** 10.1101/2024.01.31.578162

**Authors:** Rami Kersh-Mellor, Stephen H. Montgomery, Callum F. McLellan

**Affiliations:** School of Biological Sciences, University of Bristol, BS8 1TQ

**Keywords:** Lepidoptera, larvae, gregariousness, marginal predation, aposematism

## Abstract

Larval Lepidoptera gain survival advantages by aggregating, especially when combined with aposematic warning signals, yet reductions in predation risk may not be experienced equally across all group members. Hamilton’s selfish herd theory predicts that larvae which surround themselves with their group mates should be at lower risk of predation, and those on the periphery of aggregations experience the greatest risk, yet this has rarely been tested. Here, we expose aggregations of artificial ‘caterpillar’ targets to predation from free-flying, wild birds to test for marginal predation when all prey are equally accessible, and for an interaction between warning colouration and marginal predation. We find that targets nearer the centre of the aggregation survived better than peripheral targets and nearby targets isolated from the group. However, there was no difference in survival between peripheral and isolated targets. We also find that grouped targets survived better than isolated targets when both are aposematic, but not when they are non-signalling. Our data suggest that avian predators preferentially target peripheral larvae from aggregations, and that prey warning signals enhance predator avoidance of groups.

## Introduction

Understanding the evolutionary forces that favour the formation and continuation of animal groups is a key focus of behavioural ecology. A well-studied benefit of defensive grouping is the so-called ‘dilution effect’ [1-3], referring to the reduction in a given individual’s predation risk due to the lower probability that it will be singled-out by a predator. However, the survival advantages conferred by grouping are not always experienced equally across individuals in a group [e.g. 4-6]. Hamilton’s [7] ‘selfish herd’ hypothesis posits that individuals can further reduce their risk of predation by surrounding themselves with conspecifics and remaining physically close to them. By doing so, prey are said to minimise their ‘domain of danger’ (DOD), or the probability that the individual will be selected by a predator over a group mate [4,7,8]. As such, a key assertion of the selfish herd hypothesis is that an individual’s predation risk is associated with their spatial position within a group [7,8]. Peripheral individuals should be at the greatest risk, since they have the largest DOD, while those at the centre of a group should be most protected. Similarly, this theory of marginal predation [7] includes the possibility that peripheral individuals also experience greater risk if predators attack those that are nearest to them [e.g. 9]. The effect of group positioning on survival is supported by simulation models and empirical evidence [4,6,9,10]. However, aggregated species may not always behave in a way that the selfish herd hypothesis dictates [11], and predators may preferentially attack peripheral prey even when they have equal access to central individuals [5,6], calling into question the predator-prey proximity explanation of marginal predation.

Selfish herd effects and marginal predation have been studied in several vertebrate taxa [5,9,12-14], but little is known of how they might shape grouping behaviour in Lepidoptera. Gregariousness has evolved many times across larval Lepidoptera [15-17], with social behaviours varying from processionary columns, to clumped aggregations. However, few studies have investigated the existence of selfish herd-type behaviours, and their effect on survival, in gregarious caterpillars. One example comes from laboratory experiments conducted by McClure and Despland [6], who found that *Drepana arcuata* caterpillars exposed to predation by spiders, stinkbugs, and parasitoid wasps sustained fewer attacks when positioned near the centre of their aggregation compared to those on the periphery. However, while McClure and Despland’s findings suggest that group position has an effect on larval survival against invertebrate predators, testing this was not their focus, and questions remain around how other relevant predators, such as birds, might preferentially attack aggregated individuals [18]. To date, only one experiment has formally tested how avian predation risk is associated with lepidopteran larval DOD, revealing that, as hypothesised, predation risk decreases with DOD for cryptic prey in static representations of processionary columns [19]. This finding suggests that there is more to be discovered about how individual larvae might improve their survival when part of a larger aggregation.

In addition to general dilution effects, aggregated lepidopteran larvae also typically gain a considerable survival advantage through aposematic warning signals, which frequently co-evolve with social behaviour [17]. Indeed, phylogenetic analyses suggest that, in lepidopteran lineages, warning colouration may be a prerequisite for the evolution of gregariousness [16,17,20]. By aggregating with other aposematic individuals, larvae may benefit from a larger, amplified warning signal, resulting in enhanced avoidance learning by predators [21-23].

The advantages of combining grouping behaviour and aposematism are well established, yet very little is known about the potential interaction between warning signals and selfish-herd effects, and how this may shape the evolution of group behaviour. Here, using artificial models of larval prey and wild, free-flying avian predators, we investigate how warning colouration and group spatial position influence predation risk in aggregated larvae. Specifically, we test whether birds preferentially attack peripheral prey when faced with an aggregation, and whether this preference is enhanced when prey are aposematic.

## Methods

### Target details

Following established protocols [17,24], we created artificial ‘caterpillars’, or targets, to test whether larval group position and colouration have an effect on their survival when exposed to wild avian predation. Targets were either non-signalling (uniform green colouration and unpalatable) or aposematic (warning colouration and unpalatable), and grouped (aggregation of six edge and six inner targets) or ‘isolated’ (a single target separate from the aggregation) (Figure 1A). Targets consisted of an unpalatable (soaked in 2.5% Bitrex solution) mealworm inserted into a waterproof paper tube (see Supplementary methods for additional details). By making the mealworms unpalatable, we aimed to reduce repeat predation attempts from a given platform, making predators’ first choice easier to determine. A total of 13 targets were attached to a platform to create one replicate; 12 targets were positioned in a group and one ‘isolated’ target was sat separately at the opposite edge of the platform to the group, ∼4cm away (Figure 1A). We included the grouped and isolated targets on a single platform so that all would be equally visible and accessible to an approaching predator. For the main analyses all six inner targets of the group were treated as one level, but for some additional analyses the two targets at the very centre of the group were considered as a separate level: ‘centre’ (Figure 1A).

**Figure 1.**
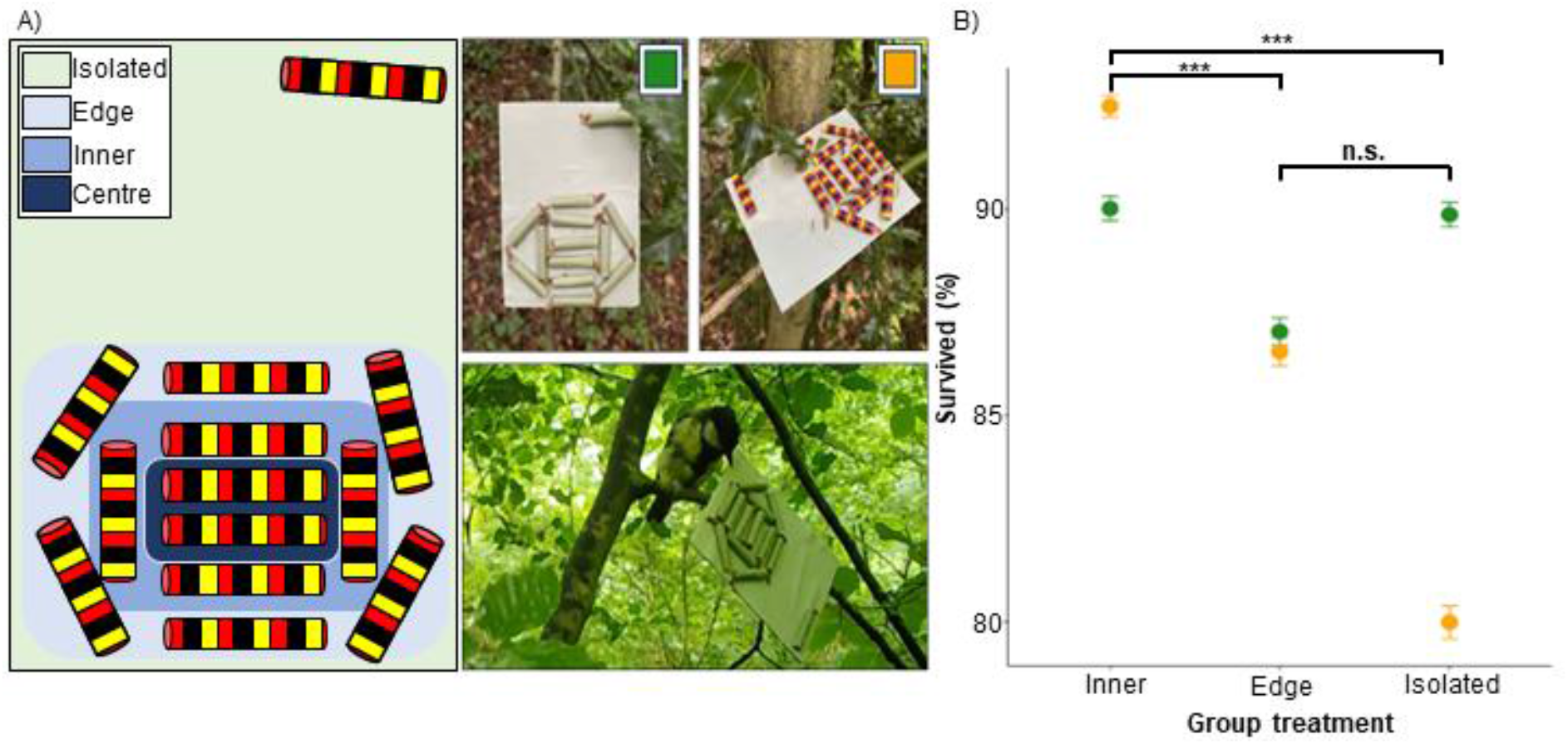
A) Examples of target layout on platforms. Left panel: coloured areas representing which targets were considered under which treatment levels in analyses. Right panel: (clockwise from top-left) platform containing non-signalling targets, platform containing aposematic targets, camera trap image of a typical avian predator attacking non-signalling targets (photographs by R. Kersh-Mellor). B) Target survival across the study, separated by colour (green points = non-signalling, orange points = aposematic) and group treatment. Whiskers represent standard deviation. Inner targets survived better than edge and isolated targets overall, whereas there was no difference in survival between edge and isolated targets.

### Survival protocol

The experiment was carried out during July 2023 around Leigh Woods (51.46259, -2.64041) in Bristol, United Kingdom. We selected sites that had abundant foliage to attach the platforms and were a reasonable distance from footpaths to avoid disturbance by humans and dogs. Platforms were checked after 4h and 24h and predation events were recorded when the mealworm was mostly or completely missing from the tube. Each experimental block was conducted at a separate site, approximately 30m away from one another, and consisted of 16 platforms, with eight replicates per treatment group. We ran a total of 10 blocks, using 2,080 targets across the study. Additionally, we used a Browning Strike Force Pro XD Nature Trail camera to record videos of avian predation (supplementary video). The camera was attached to a tree with one of the platforms in-frame. We repeated this across 10 independent platforms. These data were analysed separately from the main field data.

### Quantitative analyses

All analyses were performed in R [25]. We used the glmer function within ‘lme4’, with binomial error and a logit link function [26], to analyse treatment interactions and their effect on target survival. Our two main models tested the interaction between target colour (two levels: non-signalling and aposematic) and group position (two levels: inner and edge) and its effect on target survival, and the interaction between target colour and platform position (two levels: group and solitary) and its effect on target survival. We also tested the interaction between target colour and group position as three levels (centre, inner and edge), using the ‘emmeans’ package [27] to compare between levels. Chi-squared tests were used to analyse all predation events observed in trail camera footage.

## Results

Between the grouped targets, group position had a significant effect on survival, with inner targets more likely to survive than edge targets (X^2^ = 34.056, d.f. = 1, *P* < 0.001; Figure 1B). Centre targets also survived better than edge targets (z = 4.529, P < 0.001), but not inner, non-central targets (z = 0.788, P = 0.710). The interaction between target colour and group position was not significant (X^2^ = 0.965, d.f. = 1, *P* = 0.326). Between grouped and isolated targets, grouped targets survived better than isolated targets overall (X^2^ = 4.514, d.f. = 1, *P* = 0.034). Between grouped target levels, inner targets survived better than isolated targets (z = 4.608, *P* < 0.001), but there was no difference in survival between edge and isolated targets (z = 0.644, *P* = 0.796; Figure 1B). Furthermore, the interaction between target colour and platform position was significant (X^2^ = 7.726, d.f. = 1, *P* = 0.005), where grouped aposematic targets survived better than isolated aposematic targets (X^2^ = 10.89, d.f. = 1, P < 0.001). However, there was no difference in survival between grouped and isolated non-signalling targets (X^2^ = 0.943, d.f. = 1, *P* = 0.332).

## Discussion

Little is known of the potential influence group positioning has on the survival of aggregated larvae, and less still is known of the additional protection aposematism may afford prey based on, or possibly in spite of, their group position. Our results indicate that grouping with a chemical defence reduces predation, but only for targets nearer the centre of an aggregation. Our data also reveal that grouped aposematic prey are less likely to be predated than isolated aposematic prey which neighbour the group, but this is not the case for equally unpalatable non-signalling prey, suggesting birds’ avoidance of warningly coloured prey is enhanced when prey are aggregated, even when isolated larvae are in close proximity.

Our data provide new evidence in support of the theory that group positioning and proximity to others results in differential predation risk for aggregated individuals [7]. We found that targets nearer the centre of an aggregation, with smaller DODs, were more likely to survive than those on the periphery, despite all being equally accessible to their avian predators. This suggests that, when faced with aggregated prey, birds preferentially attack peripheral individuals, even when prey have no warning signals or obvious defences. However, our data also show that predation risk does not decrease linearly with proximity to the centre of the aggregation, and only varies based on whether prey are peripherally positioned or not. This marginal predation of our targets may suggest an innate preference of predators [5], which could evolve if peripheral prey are easier to extract, such as prey that are spiny or otherwise defended, or hungrier and therefore less vigilant [e.g. 28]. While it is possible that avian predators target peripheral larvae from ‘2-dimensional’ aggregations [5] because they are closest to where they are perched, given the targets are attacked from above, this is unlikely to explain the marginal predation observed in our data. In addition, camera trap footage (see Supplementary material) reveals that passerine predators were able to easily access both peripheral and central larvae, suggesting there was no physical barrier to targeting all individuals equally.

Our data also suggest that aposematic prey survive better when aggregated compared to nearby, isolated individuals. While this finding has an intuitive explanation when considering gregarious and solitary prey, as covered in previous literature [e.g. 17], here, all targets on a given platform were equally visible and accessible, which suggests that predators were preferentially selecting isolated over grouped prey. Importantly, there is no similar effect between non-signalling, but equally unpalatable, prey. This may indicate that, in addition to dilution, prey warning signals enhance protection against predation when they are aggregated, but only when they are visibly part of the group. Aggregation of aposematic prey significantly increases the efficacy of their warning signals in deterring predator attacks [21,22,29-31]. This effect is most likely explained by predators having a greater aversion towards larger, enhanced signals, as has been shown for both singular [32,33] and grouped prey [21,29-31]. Another explanation for our findings may be that isolated prey are easier for predators to target than any grouped individual, given that prey groups present a greater load of sensory information, which is harder for predators to process [34-36]. It may be that our aposematic target’s striped pattern somehow enhanced this confusion effect, perhaps by adding a greater number of visual elements for predators to process.

In summary, our results provide new insights into the variable predation pressures faced by gregarious larval lepidopterans. That avian predators preferentially attack the peripheral prey of a 2-dimensional group suggests that there may be additional benefits to larval grouping behaviour that are yet to be explored. As such, studies using smaller prey and larger groups may be required to explore predator preferences in finer detail.

## Acknowledgements

We are grateful to Ryan Palmer for providing the lead author with the opportunity to conduct this study. Thanks also to Leigh Woods Lead Ranger, Darren Mait for allowing us to work at Leigh Woods and his assistance in gaining us permission to do so.

## Data accessibility

Data and supplementary video uploaded to Zenodo online repository, DOI: 10.5281/zenodo.10599414

