## Supplemental material for "Selfish herd effects in aggregated caterpillars and their interaction with warning signals"

### Supplementary methods

*Target details*

Targets were created by rolling waterproof paper (Rite-in-the-Rain, J.L. Darling LLC, Tacoma, WA, USA) into cylinders (22mm length, 3.4mm diameter). All targets across the study were made unpalatable by soaking dried mealworms (Wilko dried mealworms, JK House, Notts, United Kingdom) in 2.5% Bitrex aqueous solution (5 g denatonium benzoate (Merck, Darmstadt, Germany) dissolved in 200 ml water) for at least 12h. Mealworms were inserted into paper tubes with their head and thorax protruding from the opening, and secured in place with a pinch of dough (flour, warm water, vegetable oil).

One target colour treatment (non-signalling or aposematic) was used per platform. All group target mealworms were accessible to birds, confirmed by camera trap footage (supplemental video). The platforms were made of a 80mm x 100mm rectangular piece of card (1500 micron craft board, House of Card and Paper, Amazon.co.uk) with waterproof paper glued to either side. The platform background colour was a lighter shade of green to the non-signalling targets to increase detectability of the targets. A single clothes peg (clear mini pegs, WDAFLG, Amazon.co.uk) was stuck to the underside of each platform and used to attach platforms to foliage.

*Survival protocol*

Platforms holding aposematic and non-signalling targets were placed in a random order, decided using a random number generator, at least 5m away from each other. We attached platforms to a variety of vegetation (e.g. brambles, holly, tree branches) at different levels, ranging from approximately 0.2-2m in height. Targets that showed signs of non-avian predation were treated as missing data and not included in the final analyses, for example, targets predated by mice were easily recognisable since the tubes were shredded and droppings were present on the platform (Figure S1).

Two different methods were used to record predation events observed in trail camera footage. First, all attacks on targets were recorded, even if the mealworm was absent from the tube. Then, we only recorded attacks where a mealworm was removed.

*Quantitative analyses*

For analysis of the two main models, we followed a stepwise model selection process, starting with the highest-order interaction term between target treatments (colour and group position or colour and platform position) as fixed effects, and Block and platform ID as random effects. In some instances, Block was not significant, in which case we continued with only platform ID as a random effect. We removed nonsignificant interactions and tested the next highest-order interaction term with a likelihood ration test using the drop1 function, repeating this process until only significant interactions remained. Significant interaction terms were investigated by splitting the data based on one factor’s levels and continuing the model selection process on each subset of data.

**Supplementary results**

Including all initial attacks on targets observed in trail camera footage, we found that edge targets were significantly more likely to be predated than inner targets ($X^2$ = 6.4, d.f. = 1, $P$ = 0.011). However, excluding targets that had already been predated, we found no difference in predation between edge and inner targets ($X^2$ = 3.571, d.f. = 1, $P$ = 0.059).

**Supplementary figure**

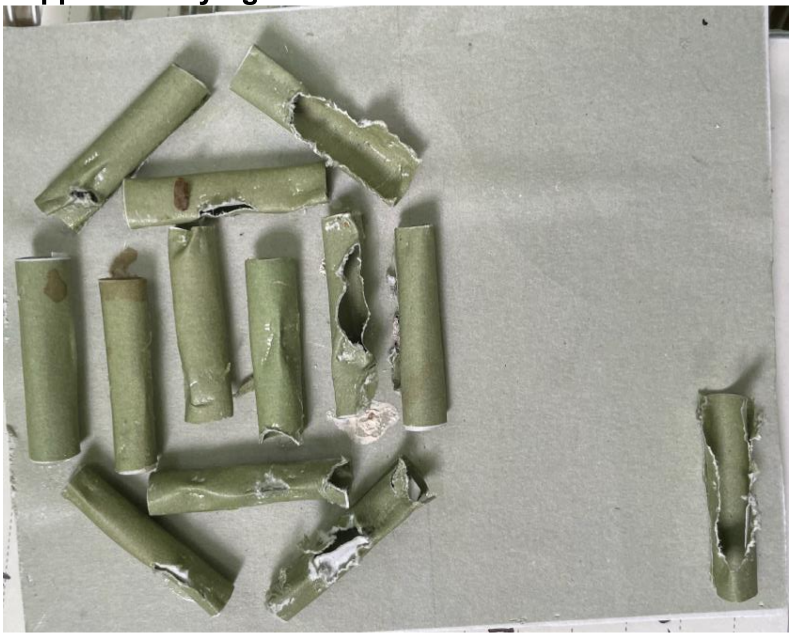

**Figure S1**. A platform of non-signalling targets showing evidence of mice predation, including droppings, shredded open targets and small paper shreds.
